# The interplay between Zfh1 and Nau orchestrates the myogenic stage-dependent expression of *Rbfox1* and its target *Stat92E* via a *mir-9a*-mediated negative feedback loop in *Drosophila melanogaster*

**DOI:** 10.1101/2024.12.02.626316

**Authors:** Amartya Mukherjee, Upendra Nongthomba

**Author notes:** Corresponding author’s.

## Abstract

RNA-binding Fox protein 1 (Rbfox1) controls gene expression at various levels: as a transcription co-factor, a splicing factor, and a regulator of mRNA stability and translation. Rbfox1 and its vertebrate orthologues have critical functions during development. *Rbfox1* mRNA, during adult myogenesis in *Drosophila,* exhibits bimodal expression, suggestive of the presence of a negative feedback loop. However, there are no known negative regulators of *Rbfox1* expression for this context. In this study, we show that the microRNA *mir-9a* functions as a repressor of *Rbfox1* expression. Furthermore, the expression of *mir-9a* is regulated by the expression of the identity transcription factor Nau. Nau and its target *mir-9a* are expressed at mid myogenesis stage, and serve to restrict the expression of *Rbfox1*, and its targets *Stat92E* and *zfh1*, to the early and the late myogenesis stages, as their ectopic expression is detrimental to early myofibrillogenesis. Zfh1, in turn, represses *mir-9a* expression, completing the feedback loop. Therefore, our findings identify the mechanism by which the temporal expression of the developmental gene *Rbfox1* and its targets are tightly regulated, essential for their functions during myogenesis.

**Summary statement:** Using *Drosophila*, we show that a microRNA-mediated negative feedback loop controls the expression of genes critical for skeletal muscle development.

## Introduction

Proper gene expression patterns during development are crucial for orchestrating the complex processes that ensure cells differentiate, migrate, and interact with each other in a coordinated manner. Gene regulation controls which genes are expressed in different cell types, allowing cells to adopt distinct fates and functions (Baralle & Giudice, 2017; Cole et al., 2023; D’Alessio et al., 2015; Tixier et al., 2010). Furthermore, gene regulation ensures that certain genes are turned on or off at specific locations and developmental stages (Bernard et al., 2003). This patterning is critical for the formation of body axes, organs, and structures (Carmena et al., 1998). Also, because development is a highly energy-intensive process, precise gene regulation ensures that cells produce only the necessary gene products for their specific functions, conserving energy and resources (Bervoets & Charlier, 2019; Breuer et al., 2019; Mori et al., 2023; Thornburg et al., 2022). Moreover, timely transcriptional/translational shutdown of specific genes is essential to prevent the overgrowth or inappropriate formation of tissues and structures (Bolondi et al., 2022; Dang et al., 2012; Posner & Laubenbacher, 2019; Wu et al., 2003). In humans, dysregulation of gene expression during development can lead to congenital disorders and developmental abnormalities (Cenik & Shilatifard, 2021; de Soysa et al., 2019; Lee & Young, 2013). Overall, precise gene regulation during development is a tightly coordinated process that involves the interplay of various regulatory elements, including transcription factors, epigenetic modifications, non-coding RNAs, and signalling pathways. This ensures that genes are expressed and silenced at the right time and in the right cells, leading to the formation of a functional and properly constituted organism.

In the model organism *Drosophila melanogaster*, optimal function comprises a set of muscles involved in flight. The dorsal longitudinal muscles (DLMs) are part of the indirect flight muscles (IFMs) in adult fruit flies, and power downward wing strokes indirectly through distortions of the thoracic walls during flight (Miller, 1950). We focused on the DLMs phenotypes because these muscles are large, and the defects are easily quantifiable, such as the total number of DLMs fascicles per hemithorax. DLMs formation and function have been extensively studied in *Drosophila* due to their relative simplicity and accessibility for experimental manipulations. Many molecular mechanisms involved in muscle development and function are highly conserved between flies and other organisms, including vertebrates (Taylor, 2006). Insights gained from studying *Drosophila* muscle development also have broader implications for understanding human muscle development and related disorders (Haigh et al., 2010; Santhoshkumar et al., 2021; Wishard et al., 2023). Moreover, the development of the DLMs can be precisely timed during the pupal stage, making it a useful system for studying the coordinated processes of muscle differentiation, migration, and attachment (Fernandes & VijayRaghavan, 1993; Fernandes et al., 1991; Gunage et al., 2017). The time-course of early DLM development encompasses the migration of myoblasts, starting at 6 hours after puparium formation (APF), to where the flight muscles develop; their surrounding, shortly thereafter, of the persistent larval oblique muscles (LOMs); their fusion with them (12-24 hours APF); and, in each hemithorax, the longitudinal splitting of the three larval muscles (LOMs) into six templates for the DLMs (between 14-28 hours APF) (Fernandes et al., 1991). The distinct patterns of gene expression suggest a precise temporal regulation corresponding to observed morphological transition points. Mutants that fail to undergo LOM splitting help identify factors which mediate splitting.

RNA-binding Fox protein 1 (Rbfox1, previously known as Ataxin-2 binding protein 1, A2bp1) affects both transcript levels and alternative splicing in muscles (Nikonova et al., 2022; Runfola et al., 2015). It contributes to normal muscle development and physiology in vertebrates (reviewed in Mukherjee & Nongthomba, 2023), but the exact mechanisms are not fully understood. In *Drosophila*, mRNA sequencing to analyse the expression profiles of genes in developing IFMs, and subsequent validation of the same at the protein level in our lab, revealed that *Rbfox1* (FlyBase Annotation Symbol: *CG32062*) expression peaks at two time-points across the developing IFMs (Nikonova et al., 2018; Spletter et al., 2018). The first peak occurs between 16 and 24 hours after puparium formation (APF), suggesting that *Rbfox1* functions during the early myogenesis stage. Indeed, our recent findings demonstrate that Rbfox1 modulates the expression of *Signal-transducer and activator of transcription protein at 92E* (*Stat92E*), the key component of the JAK/STAT signal-transduction pathway, and its effectors, and, thus, helps maintain stemness, mediates F-actin dynamics, and inhibits apoptosis in myoblasts (Mukherjee & Nongthomba, unpublished). *Rbfox1* shows a second peak occurs between 72 and 90 hours APF, suggesting that *Rbfox1* may also be involved in the late myogenesis stage, such as myofibre growth. We have previously shown that, during late myogenesis, Rbfox1 indeed affects muscle development by regulating fibre type-specific splicing and expression dynamics of numerous identity genes and structural proteins (Nikonova et al., 2022). Therefore, Rbfox1 is involved in both early and late myogenesis stages. The expression of *Rbfox1* remains significantly downregulated during mid myogenesis stage, between 30 and 72 hours APF (Nikonova et al., 2022; Spletter et al., 2018). Over-expression of *Rbfox1* using the pan-myogenic *Mef2-GAL4* driver is lethal, but its temporally and spatially restricted over-expression from 40 hours APF, using the IFM-specific *Hk^UH3^-GAL4*, results in a phenotype reminiscent of its knock-down, including torn myofibres, and thin, frayed, or torn myofibrils with short sarcomeres (Nikonova et al., 2022). This suggests that expression of *Rbfox1* during mid myogenesis stage is detrimental, and a transcriptional or post-transcriptional transition occurs during this period, allowing the muscles to undergo differentiation.

A bimodal expression pattern can be suggestive of negative feedback, a regulatory mechanism in which the downstream target of a gene inhibits its own expression. Negative feedback is a common regulatory mechanism in biology. It helps to ensure that the expression of genes is tightly controlled, and that the cells do not over-or underproduce the proteins that they need. Thus, *Rbfox1* must be modulated by regulatory molecules. Although, the transcription factors that can repress *Rbfox1* expression have not yet been identified, certain microRNAs (miRNAs) repress Rbfox1 expression. In rat hippocampal neurons, *Mir129-5p* downregulates *Rbfox1* to repress synaptic genes during homoeostatic synaptic downscaling (Rajman et al., 2017). In *Drosophila*, *mir-980*, functioning as a memory suppressor gene, represses *Rbfox1* expression to tune the excitable state of neurons (Guven-Ozkan et al., 2016). During *Drosophila* oogenesis too, *Rbfox1* levels are adjusted by *mir-980*, a stress-sensitive miRNA in this context, which increases cell viability (Kucherenko & Shcherbata, 2018). Thus, the regulation of *Rbfox1* can be complex and context-dependent, and the exact miRNAs may vary depending on the tissue or the developmental stage.

In this study, we found that a *Drosophila melanogaster* miRNA, *mir-9a*, part of an ancient miRNA family (Christodoulou et al., 2010; Fig. 1B) which includes the vertebrate *miR-9-5p*, acts as a repressor of the gene *Rbfox1*. The expression of *mir-9a* is regulated by the transcription factor Nautilus (Nau), encoded by the sole *Drosophila* orthologue of human *myogenic differentiation 1* (*MYOD1*). *Nau* expression is low during early myogenesis stage, resulting in low *mir-9a* expression. Concomitantly, Rbfox1 directly consolidates the repression of its regulator, and indirectly through Stat92E, by upregulating *Zn finger homeodomain 1* (*zfh1*), the gene product of which keeps *mir-9a* expression in check. Nau is expressed during the mid stage of myogenesis, and outcompetes Zfh1, allowing the expression of *mir-*9a, and the consequent downregulation of Rbfox1 and its targets. This is because the expression of *Rbfox1* is detrimental to early myofibrillogenesis (Nikonova et al., 2022). Later in myogenesis, the expression of *Nau* and *mir-9a* declines, which allows *Rbfox1* and its targets to be expressed again. Thus, our findings provide a novel example of how the expression of a developmental gene can be tightly controlled in a temporal manner. This may lead to a better understanding of how human *RBFOX* and *MIR9* genes may be involved in myogenesis.

**Fig. 1.**
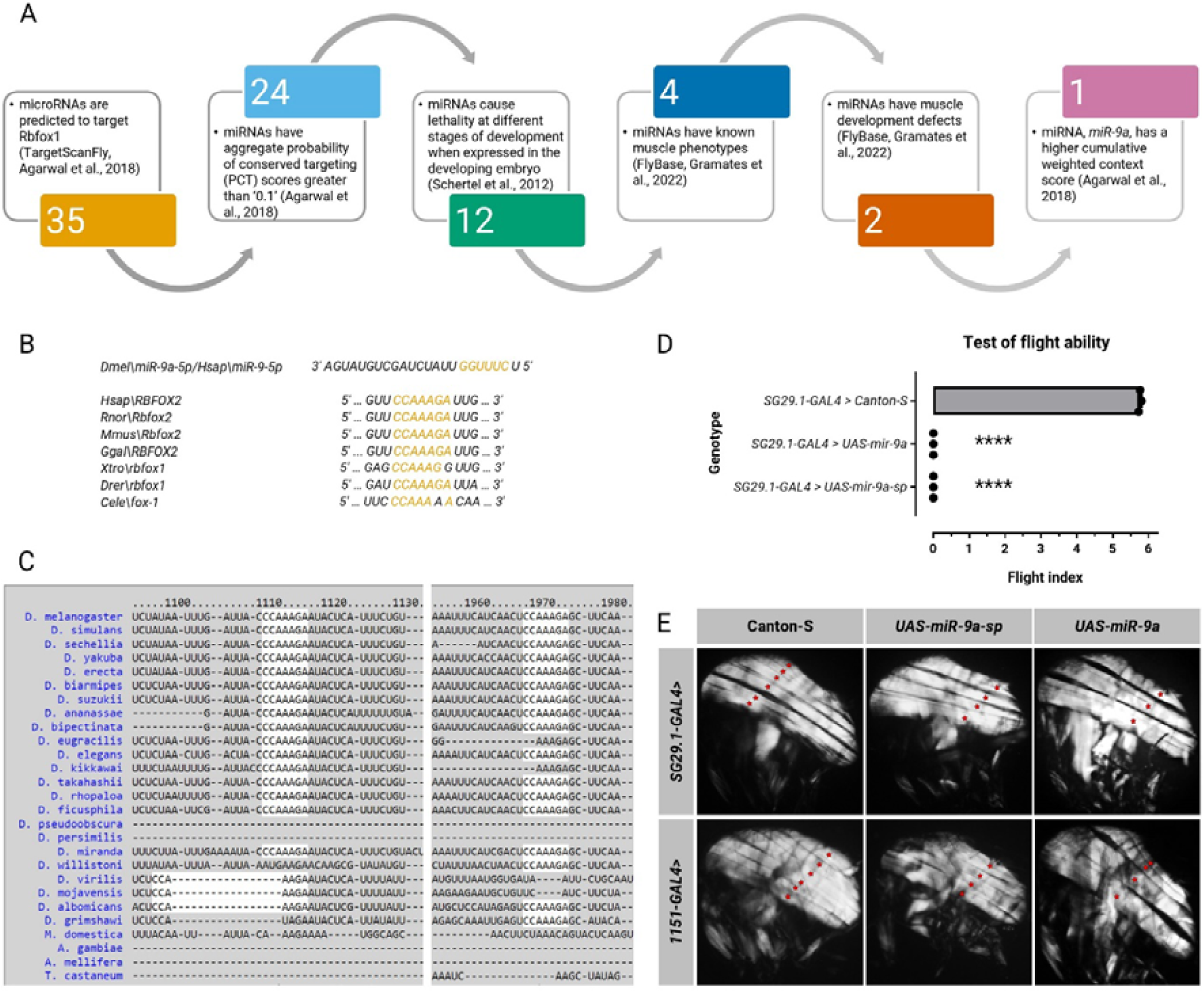
The conserved microRNA *mir-9a* negatively regulates *Rbfox1* and its targets. **(A)** Bioinformatic workflow leading to the identification of the *mir-9a* as the negative regulator of *Rbfox1.* **(B)** (above) The nucleotide sequence of *mir-9a*, and (below) its conserved 3’-UTR binding site in selected model organisms (*Hsap*, *H. sapiens*; *Rnor*, *Rattus norvegicus*; *Mmus*, *Mus musculus*; *Ggal*, *Gallus gallus*; *Xtro*, *Xenopus tropicalis*; *Drer*, *Danio rerio*; *Cele*, *C. elegans*. **(C)** The two binding sites of *mir-9a* on the *Rbfox1* 3’-UTR are conserved across drosophilids. **(D)** Quantification of flight ability after *mir-9a* mis-expression. Genotypes as noted. Significance is from paired t test (\*\*\*\**p* < 0.0001). **(E)** Polarised microscopy images of hemithorax from flies. Genotypes as noted. Red stars indicate DLM fascicles.

## Results

### Bioinformatic identification of miRNAs which putatively target Rbfox1

A study of the embryonic expression patterns of all known *Drosophila* miRNA loci found that these miRNAs are expressed in a discrete temporal and spatial manner, suggesting that they fulfil distinct roles during the development of muscle and other tissues (Aboobaker et al., 2005). However, the muscle-specific expression of these or other miRNAs in post-embryonic stages was not examined. To identify the miRNAs that participate in the regulation of the dynamic expression of *Rbfox1*, we searched for the regulatory fly miRNAs that are predicted to target *Rbfox1* using TargetScanFly (Agarwal et al., 2018) (Fig. 1A). Our search retrieved conserved sites for 35 conserved miRNA families on the *Rbfox1* 3’-UTR (Fig. 1A). To choose the candidate miRNA for our study, we decided to apply a series of criteria. A cutoff of scores greater than ‘0.1’ for the aggregate probability of conserved targeting (*P*_CT_) score (Agarwal et al., 2018), which ranks based upon the confidence that targeting is evolutionarily conserved, yielded 24 conserved miRNA families (Fig. 1A, Table S1). Next, we filtered for the miRNAs that caused lethality at different stages of development when expressed in the developing embryo by a constitutively active *Act5C-GAL4* driver (Schertel et al., 2012), and obtained such 12 miRNAs (Fig. 1A, Table S1). We then selected for miRNAs with known muscle phenotypes as per FlyBase (Gramates et al., 2022), and found four (Fig. 1A, Table S1). Of these, two had age-dependent muscle phenotypes, while the other two, *mir-9a* and *mir-92b*, had muscle development defects (Fig. 1A, Table S1). We found that the latter two microRNAs share several putative targets, all except two of which express in the developing IFMs (supplementary material Fig. S3A, Table S2). Further analysis revealed that the genes in the intersection subset are enriched for Gene Ontology terms such as ‘positive regulation of growth’, ‘regulation of mRNA metabolic process’, and ‘regulation of DNA-templated transcription’ (Table S2), strongly suggesting that *mir-9a* and the *mir-92b* converge on biological processes involved in the regulation of gene expression in the nascent myotube. We decided to focus on *miR-9a* in this study because it has two conserved sites on the *Rbfox1* 3’-UTR (Fig. 1C, Table S1), compared to the single site of *mir-92b*, and, consequently, has a higher cumulative weighted context score (Agarwal et al., 2018), a metric for the predicted repression. *mir-9a* belongs to an ancient miRNA family (Christodoulou et al., 2010), and we found that its binding site is conserved in the 3’-UTRs of *Rbfox* orthologues from *Caenorhabditis elegans* to *Homo sapiens* (Fig. 1B).

### Optimum expression of mir-9a is essential for the formation of mature DLM fascicles

Our lab has previously shown that the knock-down of *Rbfox1* driven by *αTub84B-GAL80^ts^;Mef2-GAL4* affects DLM structure and number, produces myofibre loss and sarcomere phenotypes, and the surviving adults are completely flightless (Nikonova et al., 2022). *Mef2-GAL4* drives expression in most adult somatic muscle cells (Ranganayakulu et al., 1996). In a different study, we have also observed that flies which express *UAS-mir-9a* under the control of *Hk^UH3^-GAL4* have severely disrupted DLMs: the muscles are broken and hypercontracted, and some fascicles are missing; and the sarcomeres are disorganised, and no clear Z-discs are formed (Katti et al., 2017). This results in wing posture defects and flightlessness (Katti et al., 2017). *Hk^UH3^-GAL4* drives ubiquitous expression in early pupae, becomes more pronounced in the developing IFMs by 35 hours APF, and is restricted to the IFMs from 55 hours APF (Singh et al., 2014). Therefore, at first glance, *Rbfox1* knock-down appears to phenocopy *mir-9a* over-expression.

To investigate the effect of *mir-9a* over-expression specifically during early myogenesis stage, when *Rbfox1* transcripts first peak, we decided to use *sd^SG29.1^-GAL4* (henceforth referred to as *SG29.1-GAL4*), which drives strong expression in IFM progenitor myoblasts (Roy et al., 1997; Shyamala & Chopra, 1999), and *1151-GAL4*, specific to disc-associated myoblasts (Roy & VijayRaghavan, 1997). Flies of both sexes, over-expressing *miR-9a*, exhibited a complete loss of flight ability, unlike their control counterparts (Fig. 1D). Also, unlike in the control flies, in flies over-expressing *miR-9a* there are only three DLM fascicles per hemithorax (Fig. 1E). This suggests that, when *mir-9a* expression is increased, there is no splitting of the LOM templates during the early pupal stage.

Next, we decided to assess the condition of *mir-9a* depletion. We achieved this using a transgenic line which allows *UAS*-directed expression of a *mir-9a* ‘sponge’ sequence, henceforth referred to as *UAS-mir-9a-sp*. Consequent to *mir-9a* depletion driven by *SG29.1-GAL4*, flies were rendered completely flightless (Fig. 1D). Moreover, depletion of *mir-9a* also prevents efficient splitting, as inferred from the observation that there are only four DLM fascicles per hemithorax (Fig. 1E). We noted that both the penetrance and the expressivity of this fascicular defect were lesser in comparison to that caused by *mir-9a* over-expression. Overall, these results demonstrate that a delicate balance of *mir-9a* expression is required for the splitting of the persistent LOMs into DLM templates, and, in this way, is required to regulate DLM fascicle number.

### Balanced mir-9a expression during early adult myogenesis allows the activity of its target genes

Splitting defects results from perturbed DLM biogenesis due to perturbations in the multi-step process, which involves differentiation of DLM myoblasts, their migration towards the larval templates, and myoblast fusion with these templates, as detailed above. Therefore, we investigated whether any of the gene products, which govern various aspects of this process, are targets of *miR-9a*. Bioinformatics analysis, using TargetScanFly (Agarwal et al., 2018), identified 194 potential conserved *miR-9a* targets (Fig. 2A, Table S3). Functional annotation, using PANTHER (Thomas et al., 2022), revealed that these targets are involved in a variety of cellular and developmental functions, such as signalling, RNA metabolism, cell adhesion, cytoskeletal regulation, protein degradation, apoptosis, etc. Interestingly, the largest protein class of the targets are gene-specific transcriptional regulators. Of the 48 classified putative targets which are involved in biological regulation, all, except one, are expressed during IFM development (Spletter et al., 2018) as per a transcriptomics resource (Fig. 2A, Table S3). We hypothesised that the defective splitting phenotype results due to the downregulation of a putative target, and, hence, we screened for the genes that have expression peaks at early myogenesis stage, and whose expression needs to be downregulated after myoblast fusion in pupae (Fig. 2A, Table S3). We found 16 putative target genes with such temporal expression profiles (Fig. 2A,B, Table S3).

**Fig. 2.**
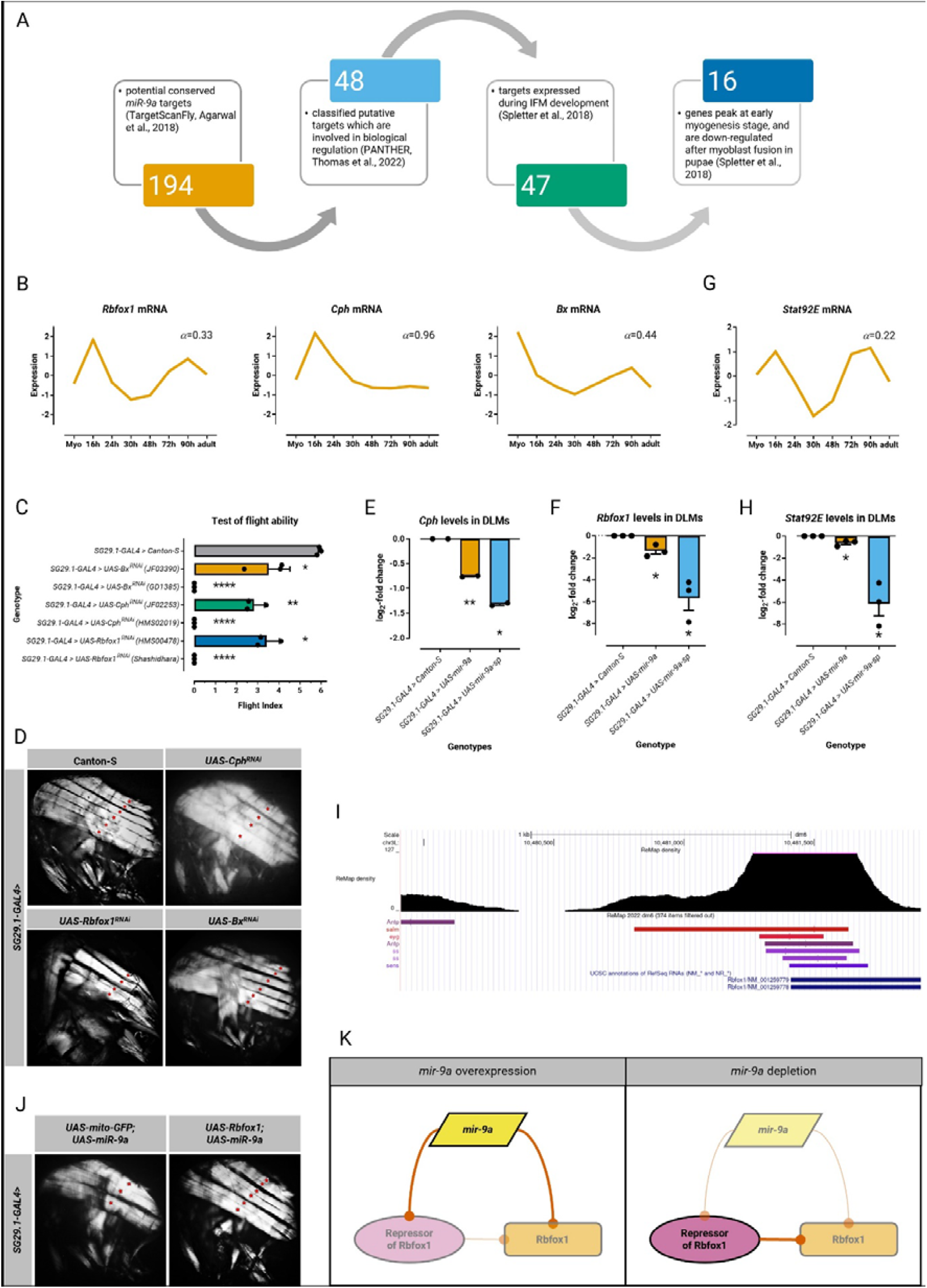
*Rbfox1* and other targets of *mir-9a* function during early adult myogenesis. **(A)** Bioinformatic workflow leading to the identification of the putative targets of *mir-9a* which may be involved in LOM splitting. Standard normal count values for **(B)** *Rbfox1*, *Cph*, and *Bx*, and **(G)** *Stat92E* from an mRNA-seq developmental time-course of wild-type IFMs (Spletter et al., 2018). *Rbfox1* and *Stat92E* have similar temporal expression profiles. **(C)** Quantification of flight ability after knock-down of putative *mir-9a* targets. Genotypes as noted. Significance is from paired t test (\**p* < 0.05; \*\**p* < 0.01; \*\*\*\**p* < 0.0001). **(D, J)** Polarised microscopy images of hemithorax from flies. Genotypes as noted. Red stars indicate DLM fascicles. Quantification of RT-qPCR data for **(E)** *Cph*, **(F)** *Rbfox1*, and **(G)** *Stat92E* transcript levels in DLMs from *mir-9a* over-expression and depletion. Significance is from paired t test (\**P* < 0.05; \*\**p* < 0.01). **(I)** View near 1 kb upstream of *Rbfox1* captured from UCSC Genome Browser assembly ID: dm6. The location of Salm, Eyg, Antp, Ss, and Sens binding sites are noted. **(K)** A model of how the over-expression and the depletion of *mir-9a* phenocopy each other. (Left) *mir-9a* over-expression during early stage of adult myogenesis, downregulates *Rbfox1* expression, but also keeps its repressor in check. (Right) *mir-9a* depletion releases the repressor of Rbfox1 from *mir-9a-*mediated inhibition, which also downregulates *Rbfox1* expression. This mechanism may buffer and help maintain the stoichiometry of Rbfox1 within the normal range during early myogenesis.

We hypothesised that the phenotypes observed upon the misexpression of *mir-9a* result due to the perturbed expression of some of such target genes. To test this, we performed ‘early’ knock-down of putative targets *Rbfox1* and *Chronophage* (*Cph*, previously known by its FlyBase annotation symbol, *CG9650*) using the *SG29.1-GAL4* driver line. Early knock-downs of *Rbfox1* and *Cph* engender a significant loss in flight ability compared to controls (Fig. 2C). Moreover, counting the number of DLM fascicles in flies with knock-downs of *Rbfox1* and *Cph* revealed that there are only three or four per hemithorax (Fig. 2D). Moreover, because *Rbfox1* has a role in myofibrillogenesis (Nikonova et al., 2022), its knock-down specifically elicits a severe hypercontraction phenotype, similar to the one reported above for *mir-9a* over-expression. Thus, the knock-downs of *Rbfox1* and *Cph*, putative targets of *mir-9a*, phenocopy its over-expression condition. As a negative control, we knocked-down *Beadex* (*Bx*), a putative target of *mir-9a* which has high expression in myoblasts from third instar larval wing discs, but is significantly downregulated by 16 hours APF (Fig. 2B). Together, these data validated our hypothesis that indeed at least some of the *mir-9a* targets are required to mediate splitting. Thus, the inherently low expression of *miR-9a* during early myogenesis stage is important for the critical roles of its targets during this stage of DLM development.

To validate the relationship between *mir-9a* and its targets, we performed real-time quantitative reverse transcription PCR (RT-qPCR), and found that *Rbfox1* and *Cph* transcripts are significantly reduced in *mir-9a* over-expression DLMs (Fig. 2E,F). Surprisingly, however, the expression of *Rbfox1* and *Cph* were found to be significantly downregulated, in fact, more so, in DLMs from flies with *mir-9a* depletion (Fig. 2E,F). Moreover, a similar trend was observed in case of *Signal-transducer and activator of transcription protein at 92E* (*Stat92E*), which shares the gene expression profile of Rbfox1 (Spletter et al., 2018) (Fig. 2G,H). Our unpublished results show that Rbfox1 directly targets *Stat92E*, the key component of the JAK/STAT signal-transduction pathway, and its effectors, and, consequently, maintains stemness, and F-actin dynamics in, and promotes survival of myoblasts (Mukherjee & Nongthomba, unpublished).

Although these findings appear to be counter-intuitive at first glance, they recall, and possibly explain, the observation that both the over-expression and the depletion of *mir-9a* elicit similar phenotypes. As proposed earlier, a balanced expression of *mir-9a* is required to generate the stereotypical number of DLM fascicles. We speculate that, during adult myogenesis, *mir-9a* buffers the expression of both Rbfox1 and its repressor(s). This robust mechanism may help maintain the optimum stoichiometry of Rbfox1 during various stages of adult myogenesis. In our previous work we have demonstrated how both *Rbfox1* knock-down and *Rbfox1* over-expression are similarly catastrophic to muscle development (Nikonova et al., 2022). We surmise that, while *mir-9a* over-expression downregulates both *Rbfox1* and its repressor, *mir-9a* depletion releases the repressor of *Rbfox1* from *mir-9a*-mediated inhibition, allowing it to attenuate *Rbfox1* expression (Fig. 2K).

Assuming that the hypothetical repressor of Rbfox1 affects its transcription, we pursued the physiological relevance of these regulatory interactions using bioinformatics. First, we identified the transcriptional regulators with peaks in the promoter region of *Rbfox1*, found using ReMap 2022, a manual curation of DNA-binding experiments for *Drosophila melanogaster* (Hammal et al., 2022) (Table S4). We sifted through the 241 unique transcriptional regulators, filtering for the ones which are also putative targets of *mir-9a*, and are downregulated during the early stage of adult myogenesis (supplementary material Fig. S3B, Table S4). Our search revealed that all of the three requirements mentioned above were met by only five transcriptional regulators: the HOX-like homeobox transcription factor Antennapedia (Antp), the Paired homeobox transcription factor Eyegone (Eyg), the basic helix-loop-helix (bHLH) transcription factor Spineless (Ss), and the C2H2 zinc finger transcription factors Spalt major (Salm) and Senseless (Sens) (Fig. 2I, supplementary material Fig. S3C, Table S4). Roy et al. (1997) have reported that hemithoraces of flies with *SG29.1-GAL4*-driven over-expression of *Antp* show disruption of the IFMs, with, notably, complete absence of the DLMs. Moreover, the knock-down of *salm* under the control of *Mef2-GAL4* leads to viable but flightless animals with a reduced number of DLMs (Schönbauer et al., 2011; Schnorrer, et al., 2010). Interestingly, we have previously shown that Rbfox1 and the transcriptional repressor Salm exhibit reciprocal regulation in the IFMs and the tergal depressor of trochanter. (Nikonova et al., 2022). Further studies are required to determine these genetic interactions.

### The over-expression of Rbfox1 rescues the mir-9a over-expression phenotype

To confirm whether *Rbfox1* is a target of *mir-9a*, we reasoned that an increase in expression level of the target should rescue the phenotype of the over-expression of the negative regulator. Thus, we directed the over-expression of *Rbfox1*, in the genetic background *mir-9a* over-expression, with *SG29.1-GAL4*. We introduced a *UAS-mito-GFP* transgenic construct in the *UAS-mir-9a* genetic background to control for the GAL4 dosage. Polarised light microscopy showed that concomitant over-expression of *Rbfox1* can suppress the *mir-9a* over-expression phenotype: six DLM fascicles were observed per hemithorax (Fig. 2J). Based on these results, we infer that *Rbfox1* expression is indeed regulated by *mir-9a* during DLM biogenesis.

### Expression of mir-9a may be regulated by Nautilus

We performed a bioinformatic search for the putative transcriptional regulators of *mir-9a* using the Alibaba2 programme (Grabe, 2002), and identified that the bHLH transcription factor encoded by the *nautilus* (*nau*) gene is the only candidate which is expressed during and characterised in *Drosophila* muscle development. Its human orthologue, myogenic differentiation 1 (MYOD1), binds to E-box sequences (*CANNTG*) in the regulatory regions of target genes (Murre et al., 1989). We find E-boxes in the 445 nucleotides-long *CpG* island, and the 1 kb-long region directly upstream of *mir-9a* (Fig. 3A). Interestingly, analysis of the Chromatin immunoprecipitation followed by sequencing (ChIP-seq) data obtained from ReMap 2022 (Hammal et al., 2022) revealed that there indeed exists a Nau peak in the *CpG* island upstream of *mir-9a* (supplementary material Fig. S1A).

**Fig. 3.**
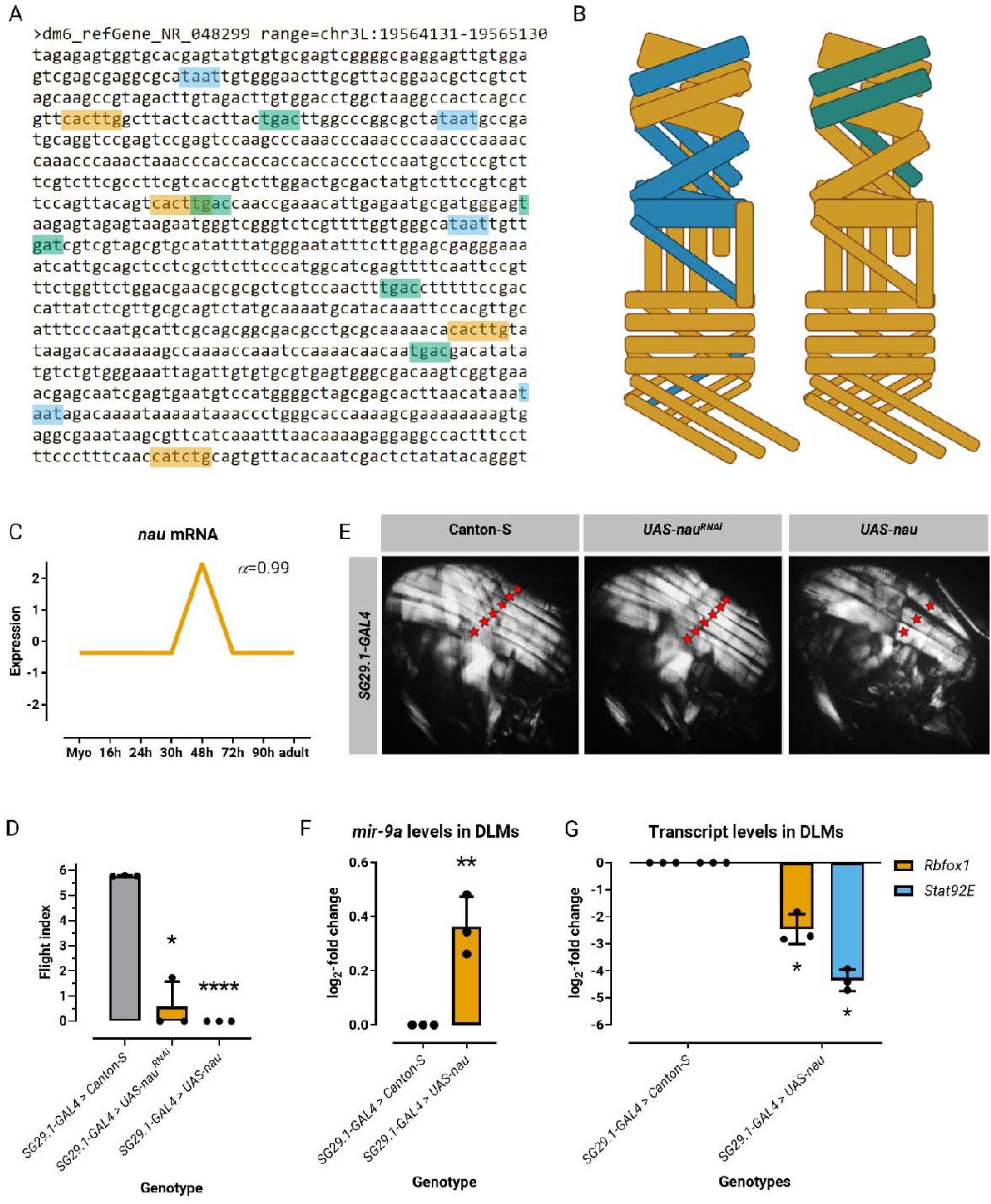
The bHLH transcription factor encoded by the identity gene *nau* regulates *mir-9a* expression during adult myogenesis. **(A)** Genomic sequence of the *mir-9a* promoter/upstream by 1000 bases retrieved from UCSC Genome Browser assembly ID: dm6. The location of Nau/Zfh1 motifs (orange), Abd-a motifs (sky blue), and Exd/Hth motifs(bluish green) are noted. **(B)** Schematic diagram depicting muscles from a hemisegment of the larva. Muscles are colour-coded to show (left) the expression pattern of the identity gene *nau*, and (right) the three larval muscles, the LOMs, that split and form muscle templates the DLMs develop from. *nau* expression incompletely overlaps with the three pairs of larval oblique muscles. **(C)** Standard normal count values for *nau* from an mRNA-seq developmental time-course of wild-type IFMs (Spletter et al., 2018). *Rbfox1* and *nau* have complementary temporal expression profiles. **(D)** Quantification of flight ability after knock-down and over-expression of *nau*. Genotypes as noted. Significance is from paired t test (\**p* < 0.05; \*\*\*\**p* < 0.0001). **(E)** Polarised microscopy images of hemithorax from flies. Genotypes as noted. Red stars indicate DLM fascicles. **(F)** Quantification of RT-qPCR data for *mir-9a* levels in DLMs from *nau* over-expression. Significance is from paired t test (\*\**p* < 0.01). **(G)** Quantification of RT-qPCR data for *Rbfox1* and *Stat92E* levels in DLMs from *nau* over-expression. Significance is from paired t test (\**p* < 0.05).

*nau* mutant embryos lack subsets of muscle fibres, most often the dorsal oblique (DO) muscle 4 and the dorsal acute (DA) muscle 3 (Keller et al., 1998). An identity gene, *nau* is expressed in larval muscles DO1, DA3, DO3-5, lateral longitudinal muscle, lateral oblique muscle, and ventral acute (VA) muscle 1 during *Drosophila* myogenesis (Dobi et al., 2015) (Fig. 3B). Interestingly, two of these larval muscle pairs, DO1 and DO3 (also known as larval oblique muscles, LOM1 and LOM3), in the second thoracic hemisegment, function as templates for the adult-specific DLM fascicles (Fernandes et al., 1991) (Fig. 3B).

We analysed the developmental mRNA-Seq dataset from IFMs (Spletter et al., 2018), and found that *nau* is expressed at low levels from the myoblast stage until 30 hours APF, peaks between 30 hours and 72 hours APF, and is downregulated again after 72 hours APF until the mature adult muscle stage (Fig. 3C). It is striking how the unimodal distribution of *nau* mRNA is complementary to the bimodal one of *Rbfox1* mRNA (Fig. 2B). Because of this apparent antagonistic interaction between *nau* and *Rbfox1*, we speculated that misexpression of *nau* may recapitulate the phenotypes due to *mir-9a* over-expression, and *Rbfox1* knock-down. Thus, we drove the over-expression and knock-down of *nau* with *SG29.1-GAL4*. Either case of misexpression of *nau* resulted in poor flight ability, especially so in flies with over-expression of *nau*, which were completely flightless (Fig. 3D). Counting the number of DLM fascicles in each hemithorax, and determined that *nau* over-expression results in a failure of the LOMs to split; whereas *nau* knock-down did not affect DLM patterning (Fig. 3E). Using RT-qPCR, we could detect increased expression of *mir-9a* in DLMs from *nau* over-expression flies (Fig. 3F). Consistent with the upregulated expression of *mir-9a*, the *Rbfox1* and *Stat92E* transcripts were found to be downregulated in DLMs from *nau* over-expression flies (Fig. 3G).

Based on these results, we infer that *mir-9a* expression is indeed regulated by Nau. Two-three E-boxes are also found in the regulatory regions of *MIR9-1*, *MIR9-2*, and *MIR9-3* as well, suggesting that this regulation may be conserved, and the expression of the human *MIR9* genes may also be controlled by MYOD1 (supplementary material Fig. S2A,B,C).

### Rbfox1 may repress mir-9a transcription via the repressive transcription factor Zfh1

The complementary expression patterns of *Rbfox1* and *nau* suggest a negative feedback loop. We questioned whether Rbfox1 exerts its repressive effect on *mir-9a* expression by downregulating *nau*. The counterparts of Rbfox proteins in *Drosophila* and vertebrates regulate alternative splicing and transcript stability by binding to *(U)GCAUG* motifs, and genes with Rbfox1 motif instances are important for muscle development (Brudno et al., 2001; Carreira-Rosario et al., 2016; Nikonova et al., 2022). We have previously performed a bioinformatic survey of muscle genes with Rbfox1 binding motifs, however, *nau* does not appear on the list of putative targets of Rbfox1 (Nikonova et al, 2022). Moreover, when we interrogated the expression of *nau* in the DLMs of flies with either knock-down or over-expression of *Rbfox1*, we did not detect any significant difference in its transcript levels compared to the control (Fig. 4A). This suggests that Rbfox1 does not control the expression of *nau*. Therefore, Rbfox1 may indirectly regulate *mir-9a* expression, via some mechanism independent of Nau.

**Fig. 4.**
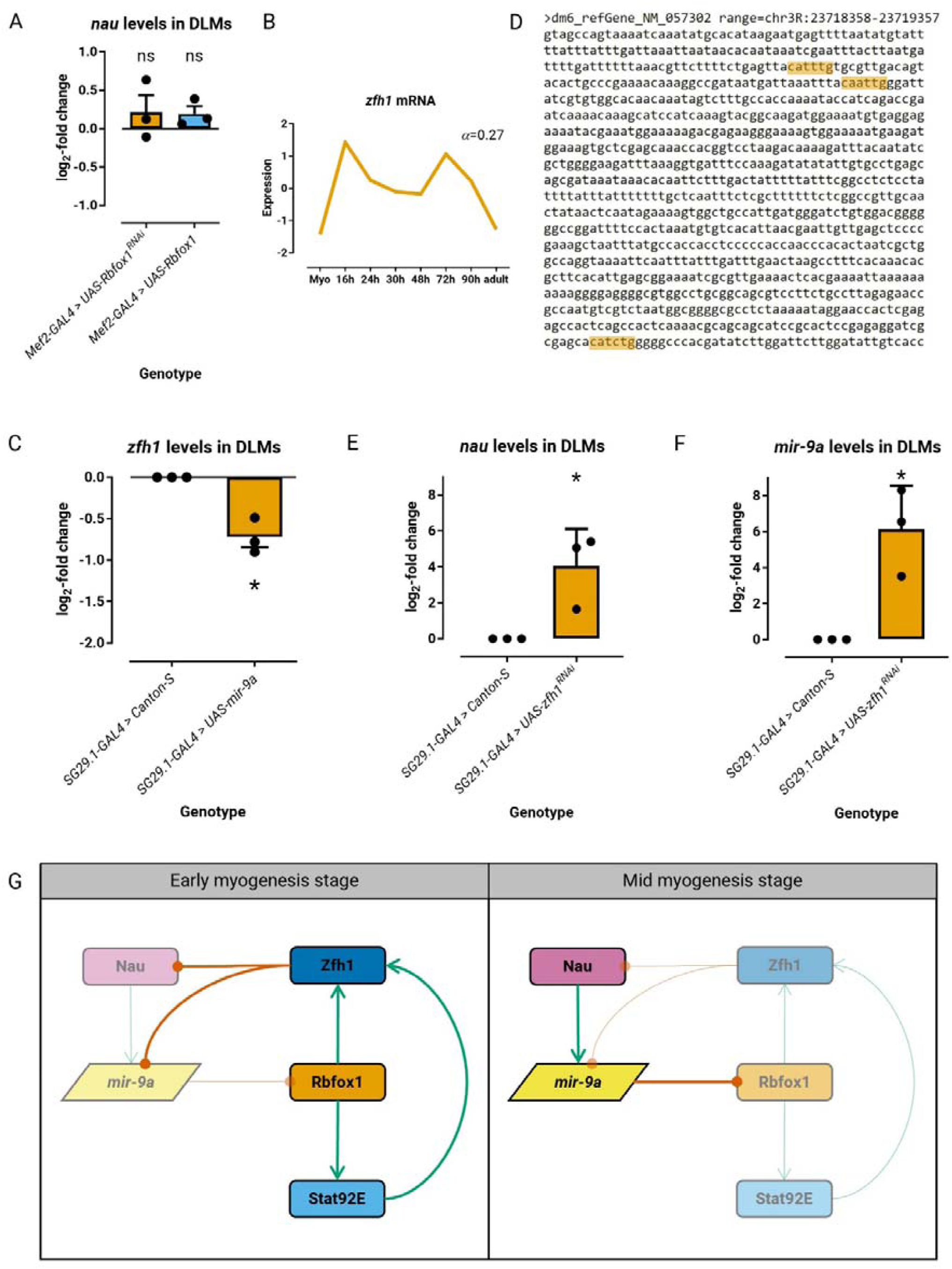
The Rbfox1 target Zfh1 may regulate *mir-9a* expression through competition with Nau. **(A)** Quantification of RT-qPCR data for *nau* transcript levels in IFMs from *Rbfox1* mis-expression. Significance is from paired t test (ns, not significant). **(B)** Standard normal count values for *zfh1* from an mRNA-seq developmental time-course of wild-type IFMs (Spletter et al., 2018). *Rbfox1* and *zfh1* have similar temporal expression profiles. **(C)** Quantification of RT-qPCR data for *zfh1* transcript levels in IFMs from *mir-9a* over-expression. Significance is from paired t test (\**p* < 0.05). **(D)** Genomic sequence of the *nau* promoter/upstream by 1000 bases retrieved from UCSC Genome Browser assembly ID: dm6. The location of Zfh1 motifs (orange) are noted. Quantification of RT-qPCR data for *nau* transcript **(E)** and *mir-9a* **(F)** levels in IFMs from *zfh1* knock-down. Significance is from paired t test (\**p* < 0.05). **(G)** A model of the negative feedback loop during adult myogenesis in *Drosophila*. During (left) early stage of adult myogenesis, the Rbfox1/Stat92E/Zfh1 module dominates, keep Nau and *mir-9a* at low expression levels. Thereafter, (right) the Nau/*mir-9a* arm gains dominance. This negative-feedback loop efficiently keeps both Rbfox1 and Zfh1 levels within the normal range during mid myogenesis.

Postigo and Dean (1997) have shown that human ZEB1 is a zinc finger homeodomain protein that represses muscle differentiation in vertebrates. Like MYOD1, ZEB1 also binds to E-boxes in muscle genes, but, unlike the former, the latter blocks transcription (Postigo & Dean, 1997). As muscle differentiation proceeds, myogenic bHLH proteins (MYOD1, MYOG, MYF5, and MYF6) accumulate to levels sufficient to displace ZEB1 from the E-boxes, which releases the repression, and allows myogenic bHLH proteins to further activate transcription (Postigo & Dean, 1997).

The *Drosophila* orthologue of ZEB1, Zn finger homeodomain 1 (Zfh1, Fortini et al., 1991) is a key downstream Stat92E effector (Leatherman & DiNardo, 2008). Moreover, *zfh1* shares the same expression profile as *Rbfox1* and *Stat92E* (Spletter et al., 2018) (Fig. 4B). This suggests that Zfh1 may repress muscle differentiation in *Drosophila* similar to how human ZEB1 does in vertebrates, and, hence, must be downregulated during mid myogenesis for the myogenic differentiation programme to proceed. We have previously shown that early IFM-specific knock-down of *zfh1* completely compromises the flight ability of flies, and causes defects in the patterning of the DLMs (Mukherjee & Nongthomba, unpublished), both phenotypes reminiscent of those caused by *mir-9a* over-expression. Our bioinformatic survey of muscle genes with Rbfox1 binding motifs (Nikonova et al, 2022) revealed *zfh1* to be a putative target of Rbfox1. Using RT-qPCR, we could detect decreased expression of *zfh1* in the DLMs from *mir-9a* over-expression flies (Fig. 4C), which suggests that *zfh1*, like its regulators, Rbfox1 and Stat92E, also lies downstream of *mir-9a*.

It is known that ZEB1 inhibits *MYOD1* expression (Siles et al., 2013; Siles et al., 2019). Indeed, we find E-boxes in the 1 kb-long region directly upstream of *nau* (Fig. 4D). We hypothesised that the knock-down of *zfh1* in IFM progenitor myoblasts would upregulate the transcription — that is, relieve Zfh1-mediated repression — of *nau.* To address this, we decided to interrogate *nau* expression in the genetic condition of knock-down of *zfh1.* Indeed, our RT-qPCR data showed that there is a significant increase in *nau* expression when *zfh1* is downregulated (Fig. 4E).

It is also known that ZEB1 represses MYOD1 transcriptional activity, and displaces MYOD1 from its DNA binding sites on target genes (Postigo & Dean, 1997; Siles et al., 2013). Therefore, it is possible that Nau shares certain E-box sequences in muscle genes, including *mir-9a*, with Zfh1. We questioned whether the knock-down of *zfh1* in IFM progenitor myoblasts would allow Nau to bind these E-boxes, and trigger the precocious expression of muscle differentiation genes such as *mir-9a*. To answer this, we decided to interrogate *mir-9a* expression in the genetic condition of knock-down of *zfh1.* Indeed, our RT-qPCR data showed that there is a significant increase in *mir-9a* expression when *zfh1* is downregulated (Fig. 4F). Moreover, our analysis of the ChIP-seq data obtained from ReMap 2022 (Hammal et al., 2022) revealed that multiple ZEB1 and MYOD1 peaks are present in overlapping fashion in the *CpG* islands and the enhancer regions upstream of *MIR9-1*, *MIR9-2*, and *MIR9-3* (supplementary material Fig. S2A,B,C). This suggests that this regulation may be conserved, and the human *mir-9* homologues may also be controlled by the competition between ZEB1 and MYOD1 during myogenesis.

## Discussion

### A novel role of mir-9a in the regulation of Rbfox1

In several biological processes, a special role is played by the microRNA-mediated feedforward loop in which a master transcription factor regulates a microRNA and, together with it, a set of target genes. From an evolutionary perspective, miRNA-mediated feedforward loops can couple fine-tuning of target protein levels with noise control, conferring precision and stability to gene expression programmes (Osella et al., 2011). In case of *miR-9a*, it promotes developmental stability by buffering *senseless* expression in the *Drosophila* proneural network (Cassidy et al., 2013; Li et al., 2006).

Various studies (reviewed in Yuva-Aydemir et al., 2011) have shown that, in vertebrates, *miR-9* is highly expressed in neural progenitor cells (NPCs), has context-dependent functions in their proliferation and migration, and regulates the generation of postmitotic neurons from NPCs. *miR-9* functions as a myogenic mirRNA too: it inhibits the proliferation and migration of smooth muscle cells by targeting *Kruppel-like transcription factor 5* (*Klf5*) (Lu et al., 2019), and the proliferation and differentiation of skeletal muscle cells by targeting *Insulin like growth factor 2 mRNA-binding protein 3* (*IGF2BP3*) (Yin et al., 2020). *miR-9a* is expressed at high levels in most epithelial cells in the *Drosophila* wing imaginal disc, and its over-expression in the wing disc can significantly suppress the development of sensory organs on the notum (Li et al., 2006). Because the myoblast cells (also known, in this context, as adepithelial cells) remain closely apposed to the epithelial surface of the notum of the wing imaginal disc, the high expression of *miR-9a* in the presumptive notum may explain the relatively low expression of *Rbfox1* mRNA and Rbfox1 protein in that region (Nikonova et al., 2022; Spletter et al., 2018; Usha & Shashidhara, 2010).

### A self-regulating mechanism mediates the dynamic expression of key myogenic genes

We propose that a negative feedback loop helps maintain the temporal expression dynamics of *Rbfox1*, which is required for proper flight muscle development. During the early adult myogenesis stage (16-30 hours APF), Rbfox1 promotes the expression of *zfh1*; Zfh1 binds to the promoter region of *miR-9a*, causing its transcriptional repression. Subsequently, the expression of *Rbfox1* and *zfh1* is downregulated, and that of *nau* is upregulated, resulting in a shift to the mid myogenesis stage (30-72 hours APF); *miR-9a* expression is derepressed. *miR-9a*, which is activated by Nau, downregulates *Rbfox1* through binding to its 3’-UTR. Again, when Nau expression declines, marking the late adult myogenesis stage (72-90 hours APF), it causes the downregulation of *miR-9a* expression. This relieves the suppression of *Rbfox1* expression caused by *miR-9a*, which accounts for the second expression peak of *Rbfox1* and its targets. Thus, Nau/*miR-9a* axis fine-tunes the level of Rbfox1 and its targets during DLM biogenesis, ensuring proper flight muscle development and differentiation.

### Rbfox1 may reinforce the regulation of mir-9a expression through transcriptional repressors additional to Zfh1

A recent study, seeking to identify how transcription factors establish cell-specific interaction networks, has demonstrated that Nau physically interacts with the three-amino-acid-loop-extension (TALE) homoeobox transcription factor encoded by *extradenticle* (*exd*) (Bischof et al., 2018). Although Exd is speculated to act as a coactivator with homoeotic proteins to activate transcription (Ryoo & Mann, 1999), in certain contexts, Exd can act as a corepressor to repress transcription with other proteins, such as Abdominal A (Abd-A) (Ryoo & Mann, 1999), and Engrailed (Alexandre & Vincent, 2003). Exd, along with the TALE homoeobox transcription factor Homothorax, confers fibrillar fate to the differentiating IFM (Bryantsev et al., 2012). The *exd*, *abd-a*, and *hth* transcripts are boosted during early myogenesis stage, and begin to be downregulated beyond 24-30 hours APF (Bryantsev et al., 2012; Spletter et al., 2018) (supplementary material Fig. S1B).

We have bioinformatically identified that both *exd* and *abd-A* contains Rbfox1 motif instances (Nikonova et al., 2022). Moreover, we have previously experimentally validated that *exd* mRNA levels were significantly decreased in Rbfox1 knock-down IFM (Nikonova et al., 2022). This suggests that Rbfox1 regulates *exd* expression. Interestingly, we find the motif instances of Abd-a and Exd, as described on the FlyFactorSurvey database (Enuameh et al., 2013), and those of Abd-a and Hth, as per the ChIP-seq data obtained from ReMap 2022 (Hammal et al., 2022), in the region upstream of *miR-9a* (Fig. 3A; supplementary material Fig. S1C). Thus, it is possible that Rbfox1 may consolidate its indirect suppression on *mir-9a* not just through Zfh1 but rather through a battery of repressive transcription factors including Exd, Abd-a, and Hth.

### A putative target of Rbfox1 may also activate mir-9a expression

We pursued, using bioinformatics, whether any direct target of Rbfox1, unlike Zfh1, may activate *mir-9a* expression. First, using ReMap 2022 (Hammal et al., 2022), we identified the transcriptional regulators with peaks in the promoter region of *mir-9a* (Table S5). Sifting through the 64 unique transcriptional regulators, we searched for the ones which are also putative targets of Rbfox1, but are upregulated during the mid stage of adult myogenesis (Table S5), and found only three that fit the criteria: the basal transcription factor encoded by *TATA binding protein* (*Tbp*), the unclassified DNA-binding domain transcription factor encoded by *lilliputian* (*lilli*), and the C2H2 zinc finger transcription factor encoded by *tramtrack* (*ttk*) (supplementary material Fig. S1D,E, Table S5). Of these, Ttk is the most promising because Ciglar et al. (2014) have reported that amorphic *ttk^D2-50^* mutant embryos exhibit a significant increase in the number of muscle founder-like cells, and decrease in the number of fusion competent cells, which, consequently, results in loss of myoblast fusion, and failure to form the normal organised muscle pattern. On the other hand, over-expression of *ttk* in the embryonic mesoderm under the control of *twi-GAL4* and *how^24B^-GAL4* was found to lead to lethality at embryonic stages due to severe defects in the specification of all three muscle types, with the somatic musculature containing very few correctly specific and differentiated muscle fibres (Ciglar et al., 2014). This suggests that Ttk may serve to initiate or reinforce the transcriptional activation of *mir-9a*, besides Nau, and may also participate in the negative feedback loop. Further studies are required to elucidate these interactions.

## Materials and Methods

### Fly strains and crosses

In accordance with the Indian Environment (Protection) Act, 1989, passed by the Government of India, the approval for work with *Drosophila* was granted by the Institutional Biosafety Committee (Ref: IBSC/IISc/UN/42/2023). Flies were reared at 25°C on standard cornmeal-agar-yeast medium. Gene nomenclature used in this report is as specified by FlyBase (http://flybase.org), unless stated otherwise. For experiments involving a wild-type control, the heterogenic strain Canton-S (FBsn0000274) was used, unless specified otherwise. The transgenic fly lines *UAS-Bx^RNAi^* (Stock # dna1385; Kairamkonda & Nongthomba, 2018), and *UAS-mir-9a* (Stock # 41138; Katti et al., 2017), *UAS-Rbfox1^RNAi^* (Stock # 32476; Nikonova et al., 2022), *UAS-Bx^RNAi^* (Stock # 29454; Kairamkonda & Nongthomba, 2018), *UAS-Cph^RNAi^* (Stock # 40852; Yamaguchi et al., 2023; and 26713; Ni et al., 2009), *UAS-Rbfox1^RNAi^* (Stock # 32476; Nikonova et al., 2022), and *UAS-nau^RNAi^* (Stock # 50607; Ni et al., 2011) were procured from the Vienna *Drosophila* RNAi Center (VDRC) and the Bloomington *Drosophila* Stock Center (BDSC), respectively. *UAS-mir-9a-sp* (Loya et al., 2009); *UAS-Rbfox1* and *UAS-Rbfox1^RNAi^* (Usha & Shashidhara, 2010); *UAS-zfh1^RNAi^* (VDRC Stock # 103205; Boukhatmi & Bray, 2018); and *UAS-nau* (Stock # FL0585; Guruharsha et al., 2011) were gifted by Prof David Van Vactor (Harvard Medical School, Boston, USA), Prof L. S. Shashidhara (Indian Institute of Science Education and Research, Pune, India), Dr Hadi Boukhatmi (IGDR, Rennes, France), and Prof K. VijayRaghavan and the Fly Facility at the National Centre for Biological Sciences (NCBS), Bangalore, India, respectively.

The GAL4/UAS system was used to drive ectopic gene expression in a tissue-specific manner (Brand & Perrimon, 1993). To preclude effects on the adult muscle precursors during embryogenesis from impacting results, we performed the knock-down experiments driven by *Mef2-GAL4* using the temporal and regional gene expressing targeting (TARGET)-inducible system, which relies on a temperature-sensitive GAL80 repressor (encoded by α*Tub84B-GAL80^ts^*) (McGuire et al., 2003). The crosses were performed at 25°C, unless indicated otherwise. Barring cases of lethality, the over-expression, the RNAi-based knock-down, and the control pupae were allowed to develop at 18°C from embryos until pupation (around 13 days), and then shifted to 29°C until eclosion as adults.

### Flight testing

Flies were aged for 2-3 days before testing for flight ability, and the assay was performed as described by Drummond et al. (1991). Flight ability was scored by releasing the flies individually from the centre of a Perspex flight chamber (henceforth referred to as ‘Sparrow’ Box, named so in honour of Prof John C. Sparrow) 40 cm high and 20 cm square with a light source at the top. Each fly was scored as horizontal (H) if it alighted at the side of the box in a zone 5 cm high and level with the release point, or up (U) if it flew up above this point. Flies dropping vertically to the bottom of the box were scored as flightless (F), with those flying between the bottom and the horizontal band scored as down (D). The flight ability data were then transformed into a flight index for each genotype, determined by using the formula described by Tohtong et al. (1995): 6 U/T + 4 H/T + 2 D/T + 0 F/T, where U, H, D, and F are the number of flies in each category of flight ability, and T is the total number of flies tested for that genotype.

### Polarised microscopy

Sample preparation for imaging the IFMs in the adults was done as described by Nongthomba and Ramachandra (1999). Flies were anaesthetised, frozen in liquid nitrogen, bisected with a razor blade, and then processed by dehydrating in ethanol series [1 hr each in 50%, 75%, 90%, and 100% ethanol (Changshu Hongsheng Fine Chemical)], and clearing in methyl salicylate (S. D. Fine Chem. Ltd., SDFCL). The head, abdomen, wing, and legs of each sagittal section were removed, using insulin syringes with ultra-fine needles, prior to mounting the hemithoraces, using DPX mountant (SDFCL). The hemithoraces were observed under an Olympus SZX12 microscope, and photographed using an Olympus C-5060 camera under polarised light optics.

### RNA isolation, and cDNA synthesis

RNA was obtained from the DLMs of 2-3 days-old flies. The flies were bisected using liquid nitrogen, transferred to 70% ethanol (Fisher Chemical), and kept at -20°C overnight. The DLMs were dissected out from the hemithoraces, and immersed in TRI Reagent (Sigma-Aldrich). The tissue was homogenised, and the total RNA was isolated by following the manufacturer’s instructions. Using the NanoDrop 1000 UV-VIS Spectrophotometer (Thermo Scientific), RNA concentration and purity were quantified. First strand cDNA was synthesised, for miRNA, using 2 µg of total RNA, and the Verso cDNA Synthesis Kit (Thermo Scientific), and, for mRNA, using 2-5 μg of total RNA, and the RevertAid First Strand cDNA Synthesis Kit (Thermo Scientific), Maxima H Minus First Strand cDNA Synthesis Kit (Thermo Scientific), the iScript Advanced cDNA Synthesis Kit for RT-qPCR (Bio-Rad), or the iScript Reverse Transcription Supermix for RT-qPCR (Bio-Rad).

### Real-time quantitative reverse transcription PCR (RT-qPCR)

Quantitative PCR was carried out using HOT FIREPol EvaGreen qPCR Supermix (5X, Solis BioDyne) or Maxima SYBR Green/ROX qPCR Master Mix (2X, Thermo Scientific). All the RT-qPCR reactions were performed at the QuantStudio 3 Real-Time PCR System (Thermo Scientific), using, in case of mRNAs, gene-specific exon-exon junction-spanning primers. The relative changes in gene expression were calculated after normalisation to the expression of a housekeeping gene, *Ribosomal protein L32* (*RpL32*, previously known as *rp49*). The expression level of *miR-9a* was determined by the RT-qPCR-based method developed by Sharbati-Tehrani et al (2008). The 2^-ΔΔC^ method was used as the relative quantification strategy the qPCR data analysis (Livak & Schmittgen, 2001). Details of the primers are provided in the table below.

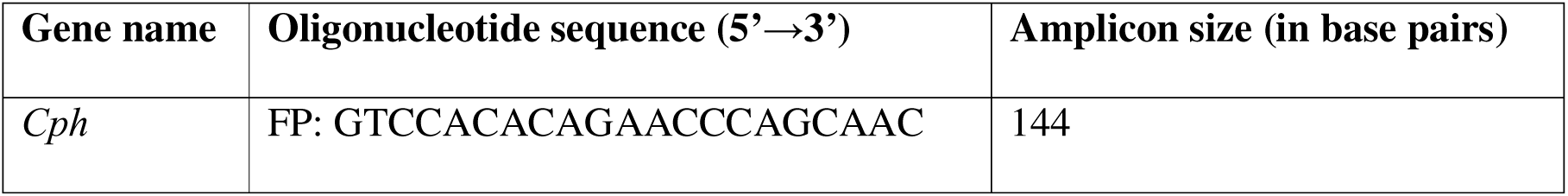
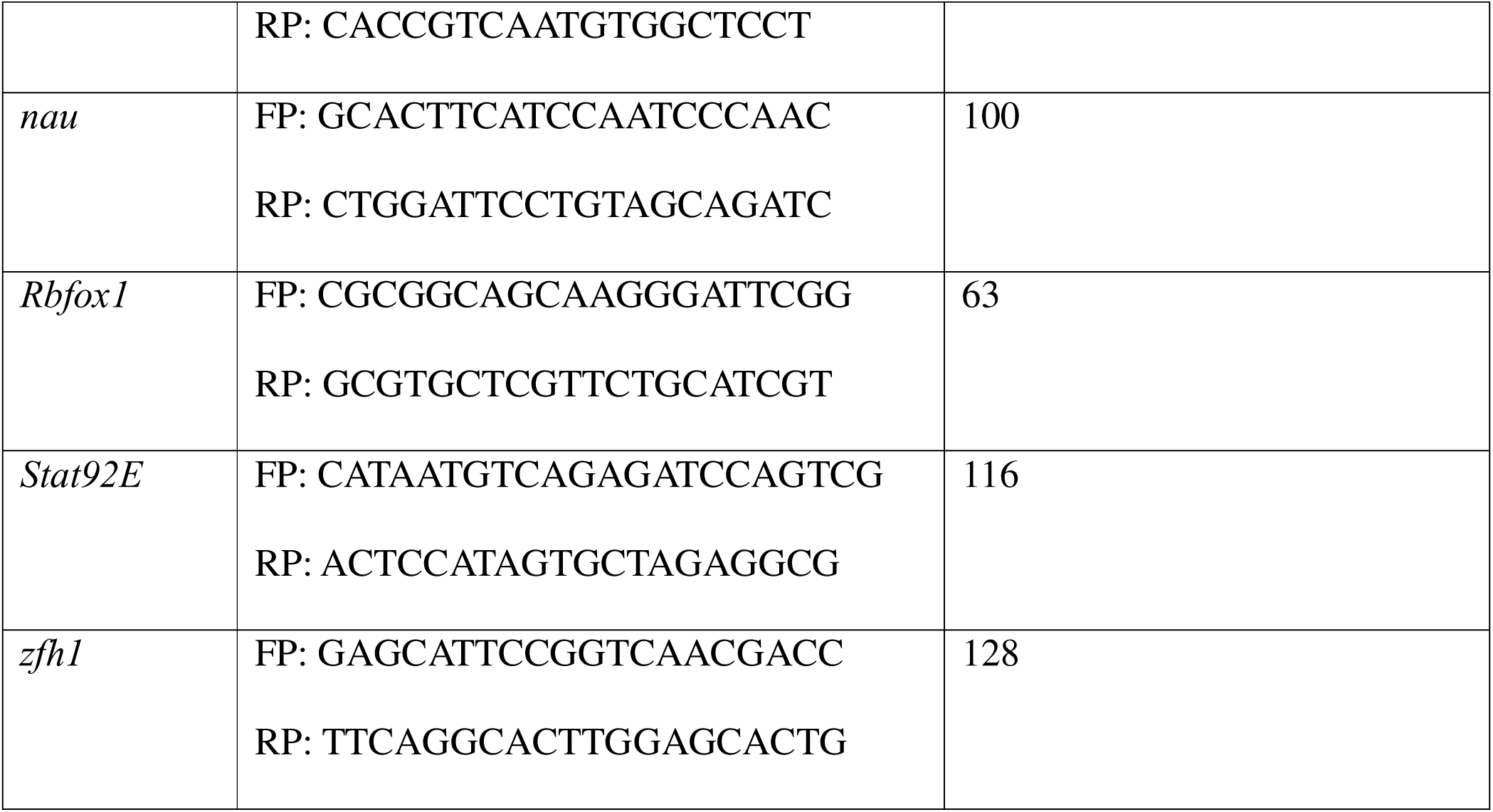

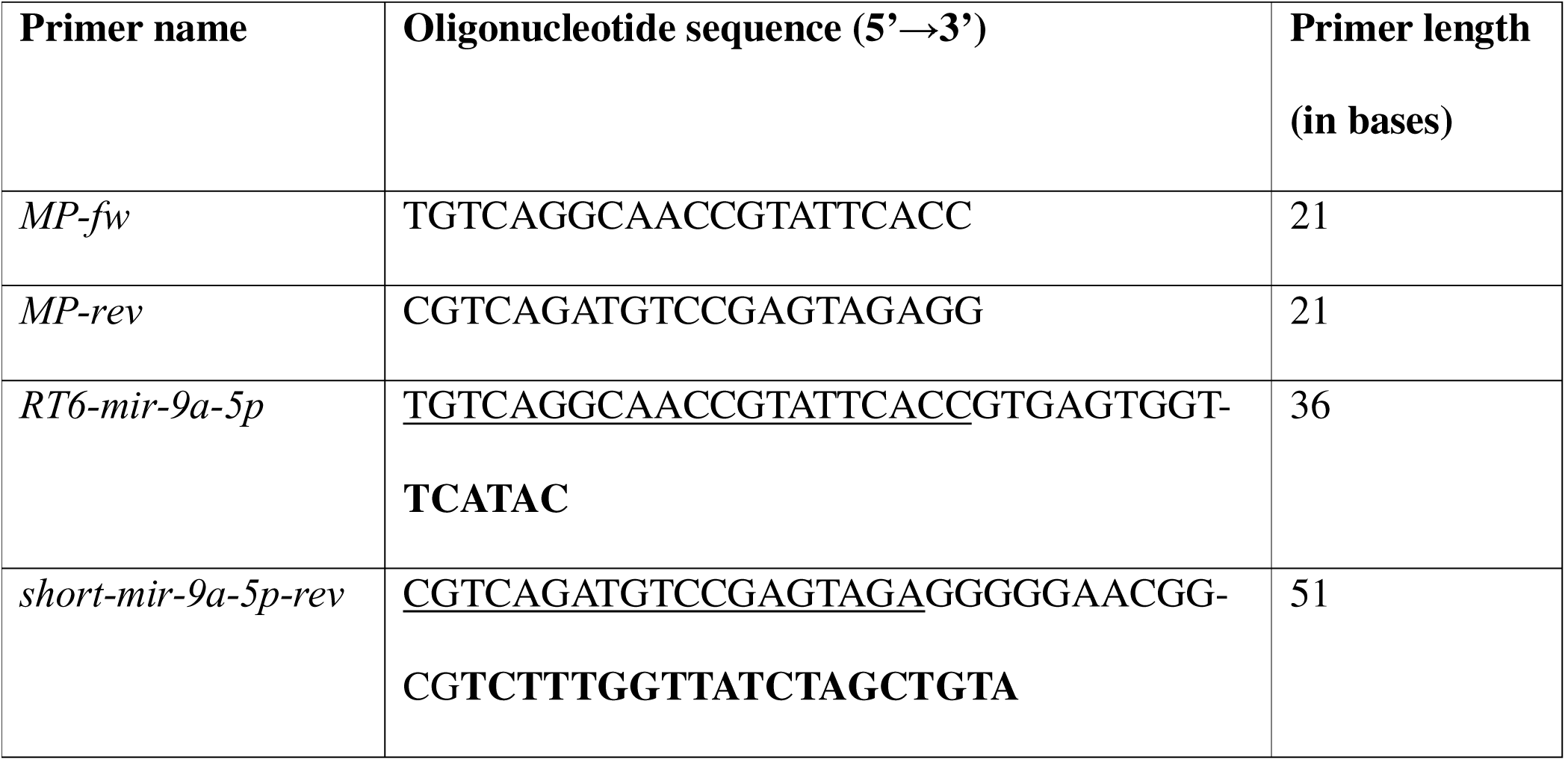

### Bioinformatic analysis

FlyBase (release FB2023_06) was used to find information on phenotypes, function, stocks, gene expression, etc. Gene Ontology term enrichment for biological process was performed using the PANTHER Overrepresentation Test (Mi et al., 2019). MicroRNA matches to *Rbfox1* 3’-UTR, and predicted fly microRNA targets of the conserved microRNA families *mir-9a-5p* and *mir-92b-3p* were retrieved using TargetScanFly 7.2 (Agarwal et al., 2018). The UCSC Table Browser tool (Karolchik et al., 2004) was used to retrieve and export ReMap 2022 (Hammal et al., 2022) track data for regions of interest.

### Statistical analysis

Statistical analysis of data shown was performed using Prism for Windows version 8.0.2 (GraphPad Software Inc., La Jolla, CA, USA). Pairwise comparisons were made using two-tailed Student’s t test. Error bars in histograms represent standard errors of means. Relevant comparisons were labelled as either significant at *p* ≤ 0.0001 (****), *p* ≤ 0.001 (***), *p* ≤ 0.01 (**) or *p* ≤ 0.05 (*) levels, or non-significant (ns) for values of *p* > 0.05.

### Data visualisation

Graphs were created using GraphPad Prism 8.0.2, and the colourblind-friendly Okabe-Ito palette (Ichihara et al., 2008). Area-proportional Venn diagrams were created using the DeepVenn web application (Hulsen, 2022). Screenshots of regions of interest from the *D. melanogaster* (Aug 2014 BDGP Rel6 + ISO1 MT/dm6) genome assembly were produced from the UCSC Genome Browser (Kent et al., 2002).

## Data availability

Fly lines are available upon request. The authors affirm that all other data necessary for confirming the conclusions of the article are present within the article, figures, tables, and in the supplementary data files.

## Acknowledgments

The authors thank Vishakha Nesari, Upasana Gupta, and Sautan Show for their helpful comments on the manuscript. They extend their sincere gratitude to BDSC, VDRC, David Van Vactor, Hadi Boukhatmi, L. S. Shashidhara, and K. VijayRaghavan and the NCBS Fly Facility for kindly providing various fly lines used in this study. The work was supported by the Indian Institute of Science Indian Institute of Science, Bangalore (IE/REDA-23-1788-18). A.M. was a recipient of the research fellowship of the Council of Scientific and Industrial Research, New Delhi [09/079(2787)/2018-EMR-I].

## Author contributions

A.M.: conceptualization (equal); funding acquisition (supporting); investigation; data curation; formal analysis; visualization; writing – original draft preparation; writing – review and editing (equal). U.N.: Supervision; conceptualization (equal); Funding Acquisition (lead); writing – review and editing (equal).

---

**Supplementary material Fig. 1.**
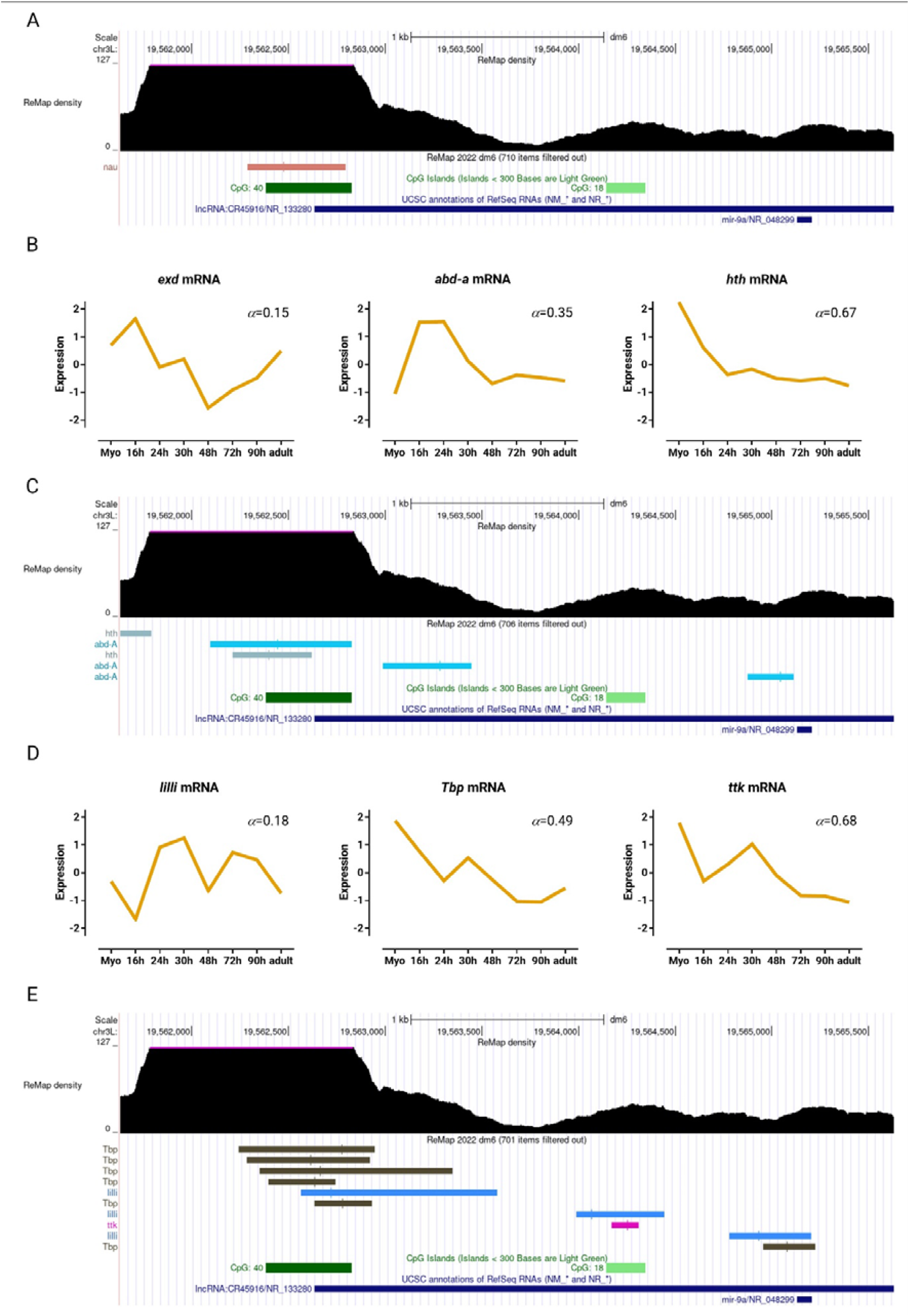
View near the *mir-9a* gene captured from UCSC Genome Browser assembly ID: dm6. The location of **(A)** Nau, **(C)** Abd-a, and Hth, and **(E)** Ttk binding sites are noted. Standard normal count values for **(B)** *exd*, *abd-a*, and *hth*, and **(D)** *lilli*, *tbp*, and *ttk* from an mRNA-seq developmental time-course of wild-type IFMs (Spletter et al., 2018).

**Supplementary material Fig. 2.**
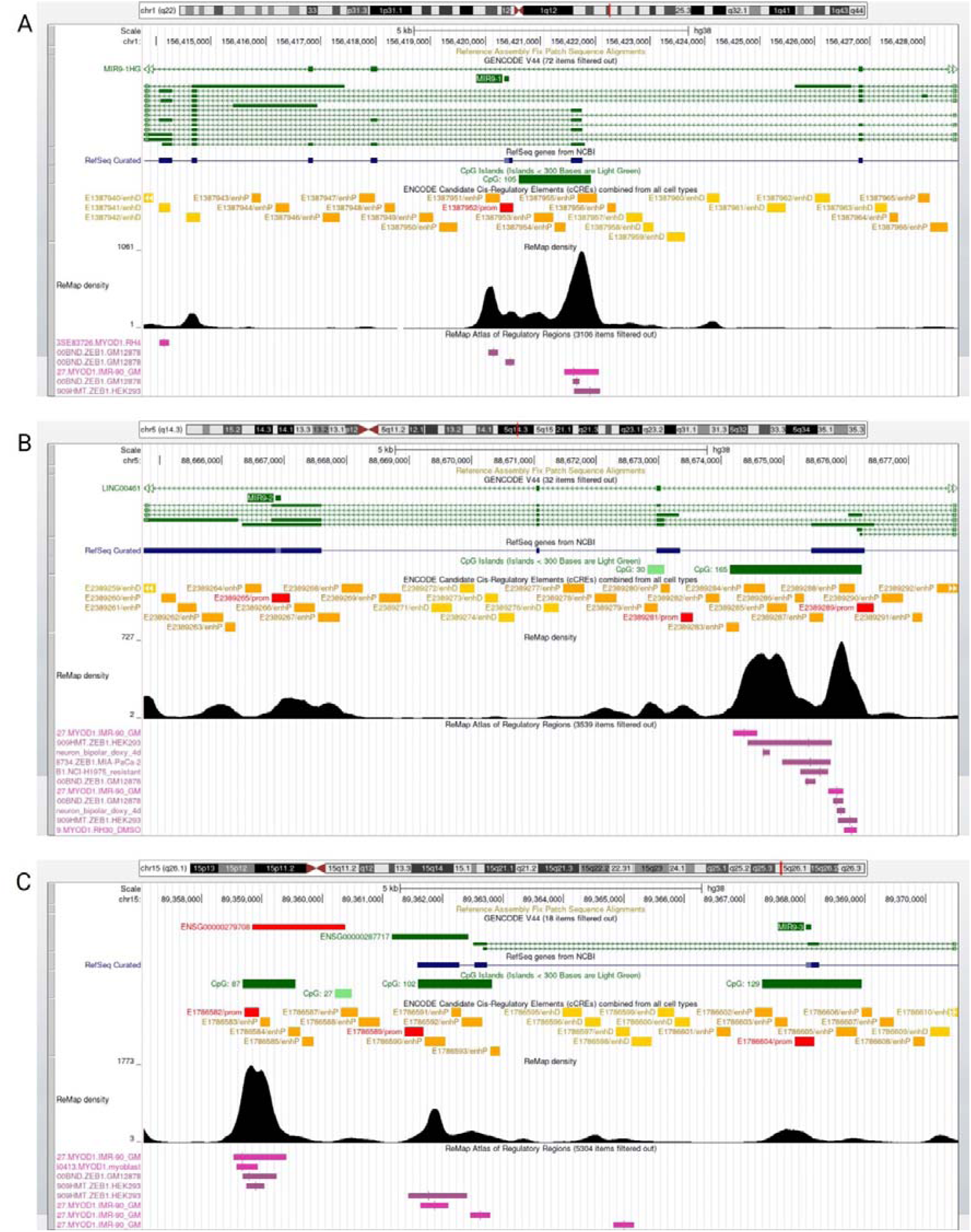
Views near the **(A)** *MIR9-1*, **(B)** *MIR9-2*, and **(C)** *MIR9-3* genes captured from UCSC Genome Browser assembly ID: dm6. The location of MYOD1 motifs (pink), and ZEB1 motifs (plum) are noted.

**Supplementary material Fig. 3.**
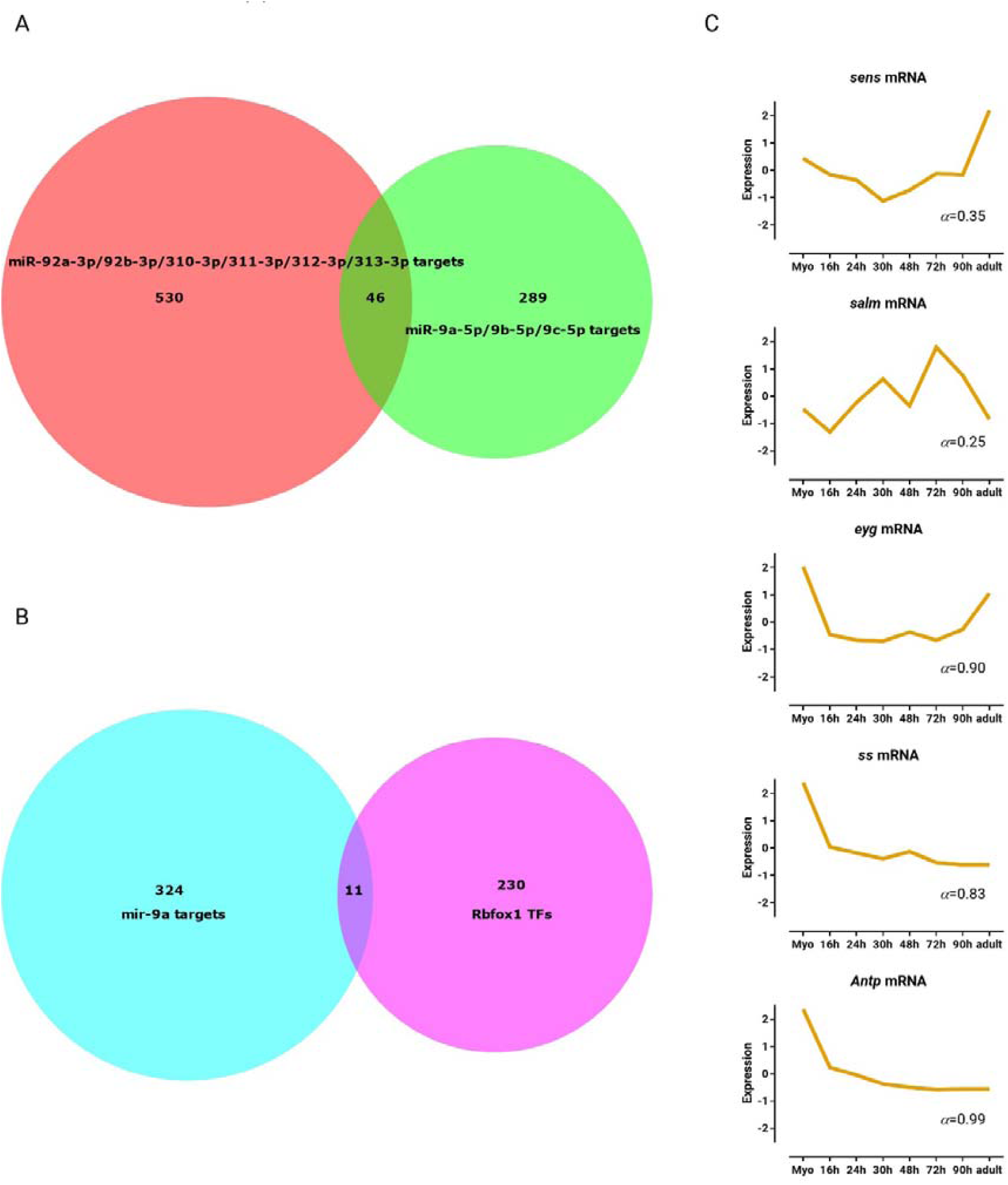
**(A)** Area proportional Venn diagrams representing the putative targets of the conserved microRNA families *mir-9a-5p* (lime) and *mir-92b-3p* (red), retrieved using TargetScanFly 7.2 (Agarwal et al., 2018). **(B)** Area proportional Venn diagrams representing the putative targets of *mir-9a-5p* (aqua), and transcriptional regulators binding to the region 1 kb upstream of *Rbfox1* (fuchsia), retrieved using TargetScanFly 7.2 (Agarwal et al., 2018) and the UCSC Table Browser tool (Karolchik et al., 2004), respectively. **(C)** Standard normal count values for *sens*, *salm*, *eyg*, *ss*, and *Antp* from an mRNA-seq developmental time-course of wild-type IFMs (Spletter et al., 2018).

