## Supplementary figures and images for "The interplay between Zfh1 and Nau orchestrates the myogenic stage-dependent expression of *Rbfox1* and its target *Stat92E* via a *mir-9a*-mediated negative feedback loop in *Drosophila melanogaster*"

### Supplementary material Fig. S1

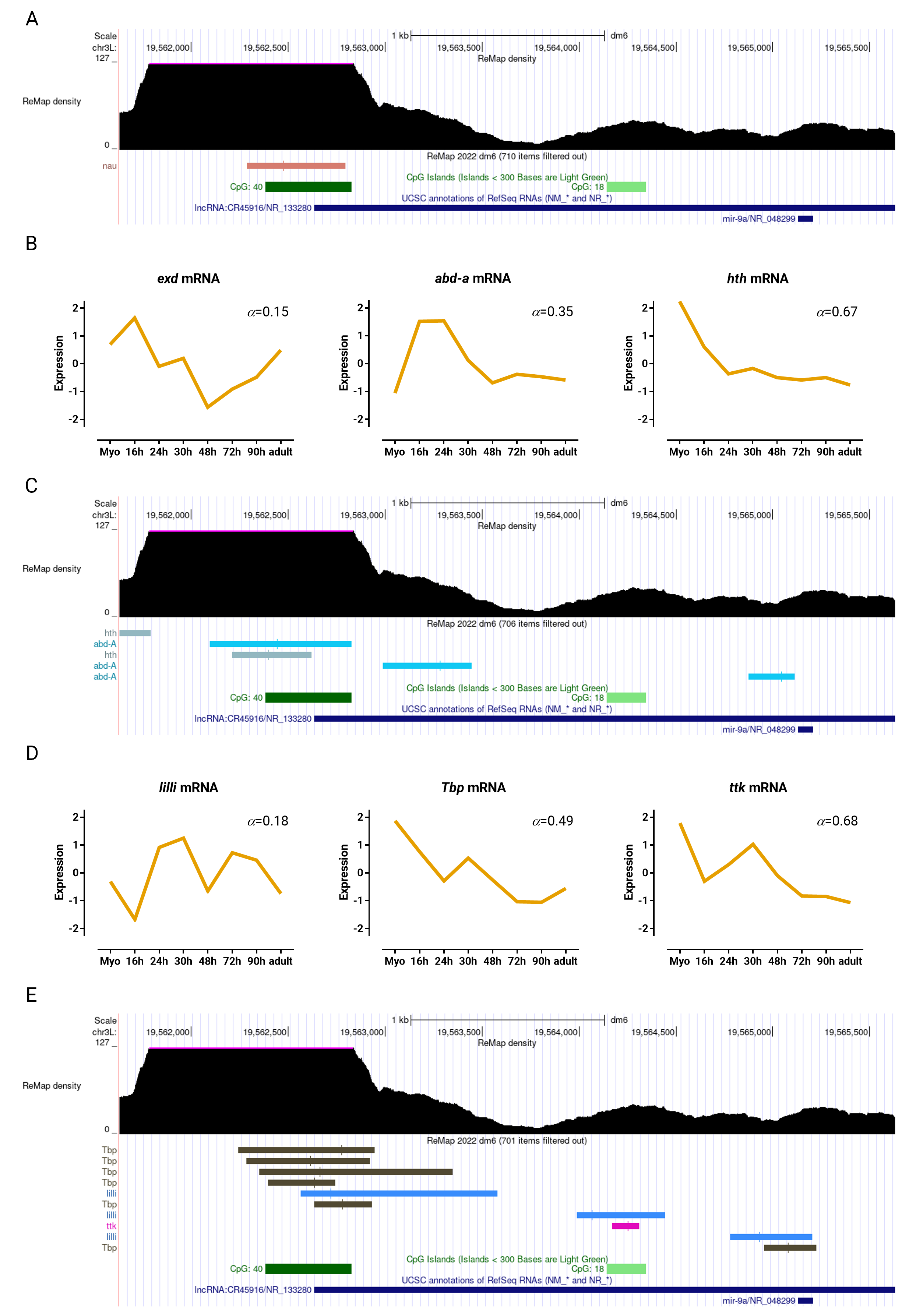

### Supplementary material Fig. S2

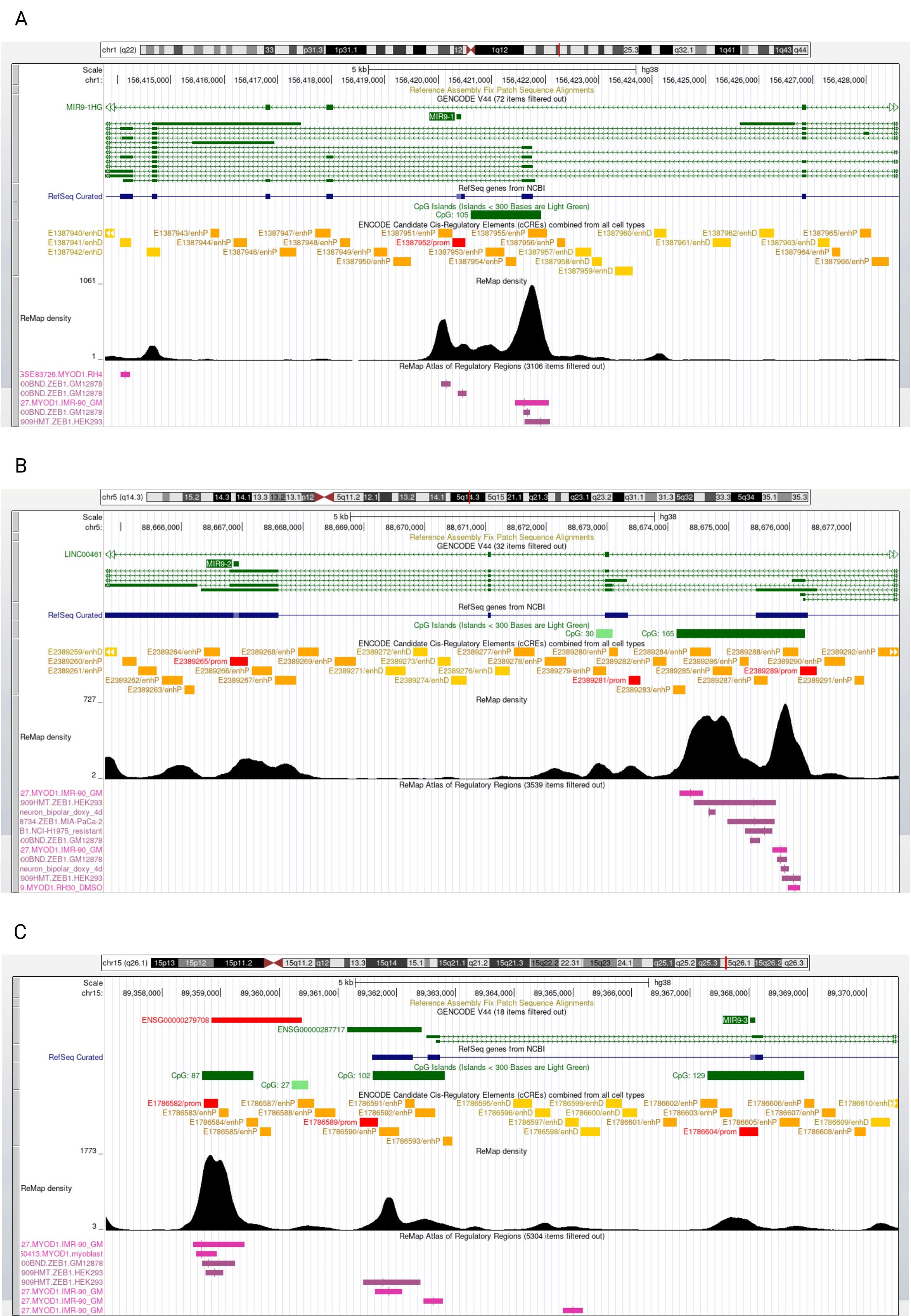

### Supplementary material Fig. S3

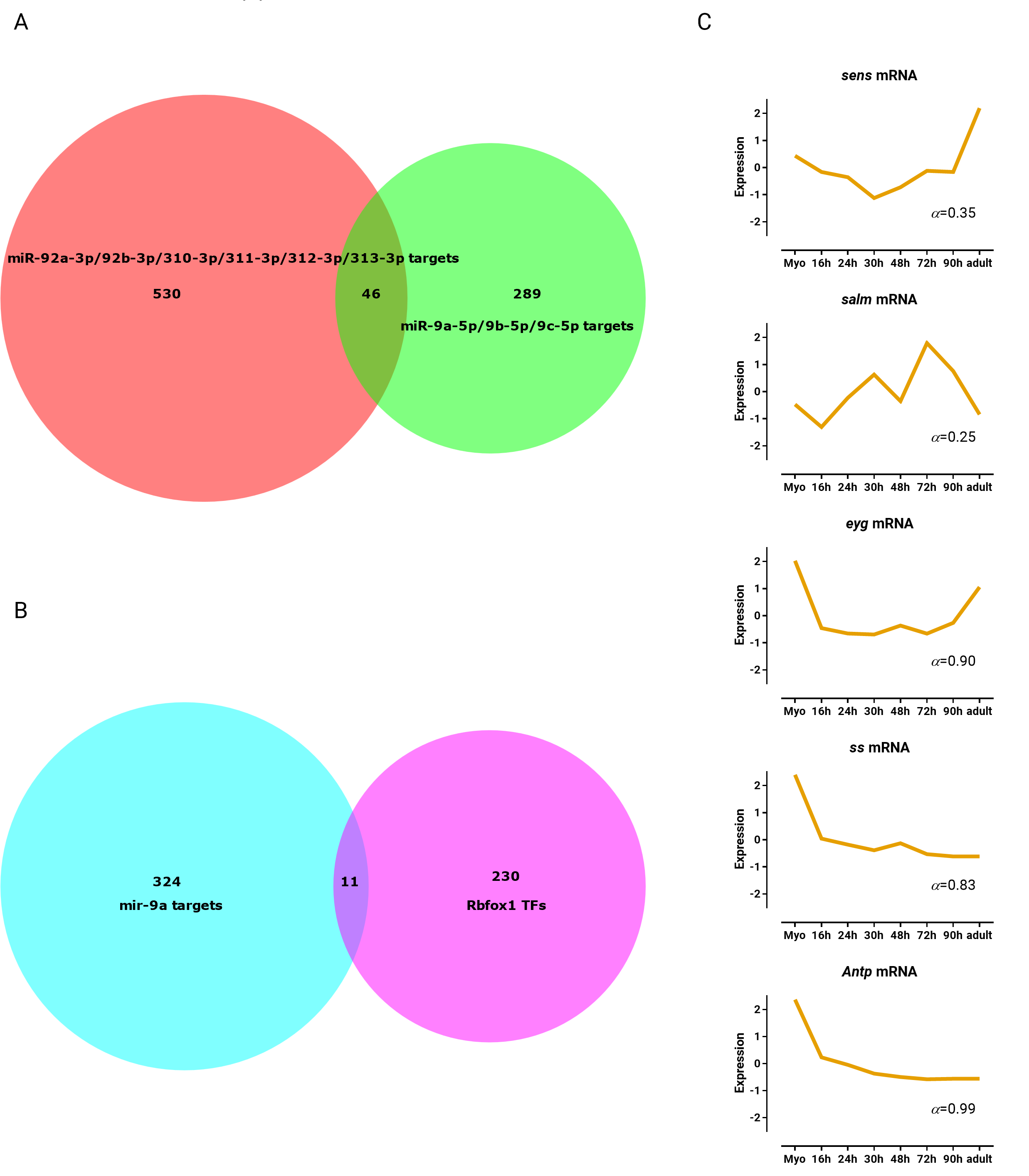
